# Responses of Soil Bacterial Community and Its Resistome to Short-term Exposure to Macrolide Antibiotic Macrolactin A: Metagenomic analysis

**DOI:** 10.1101/2025.04.29.651243

**Authors:** Darya V. Poshvina, Alexander S. Balkin, Diana S. Dilbaryan, Alexey S. Vasilchenko

## Abstract

An important aspect of studying potential antibacterial biopreparations for crop protection is determining their potential negative impacts on the environment. Plant-associated *Bacillus velezensis* produce macrolactin A (McA), which determine effectiveness of this bacterial species against numerous human and plant pathogens. However, the effects of McA on the soil microbiome and the selection of specific antibiotic resistance genes (ARGs) among soil bacteria remain unknown. In this study, we use high-throughput sequencing-based metagenomic methods to investigate the differences in structure of the soil bacterial community and the abundance and diversity of ARGs in both McA-treated and untreated samples. The presence of high (10 mg per kg soil) and low (1 mg per kg soil) concentrations of McA induced changes in soil bacterial populations as shown by taxonomic analysis. The relative abundance of Alphaproteobacteria and Betaproteobacteria significantly increased under the McA treatments, while the relative abundance of Thermoleophilia, Rubrobacteria, Planctomycetia and Acidimicrobiia decreased.

The ARG profiling results showed that both low and high doses of McA affected ARGs representation in the community. At the same time, a low dose of McA altered the representation of a larger number of ARGs (7 genes) compared to a high dose (3 genes). Overall, exposure to McA consistently altered the abundance of genes associated with resistance to elfamycin, glycopeptide, fluoroquinolone, rifampicin, and macrolide. Correlation analysis identified 185 relationships between 52 antibiotic resistance genes (ARGs) and 34 bacterial genera. Among these bacteria, *Streptomyces, Baekduia*, and *Capillimicrobium* were predicted to carry the most diverse ARGs. By assembling and annotating bacterial genomes, we identified the true hosts of ARGs. Chloroflexota were the most prevalent phylum harboring ARGs. Furthermore, profiling the soil microbiome’s metabolic potential under low-dose McA revealed increased abundance of genes associated with signaling, chemotaxis, and broad-substrate drug efflux.

The collectively obtained data significantly expands the understanding of the functional role of McA in the ecology of *Bacillus velezensis*, and at the same time provides an assessment of environmental risks associated with the use of biopreparations containing metabolites or living cells of this species’ bacteria.

## 1. Introduction

As the world’s population grows and the demand for high-quality, affordable food increases, the importance of organizing effective crop protection against various stresses is of global significance (Ali N. et al., 2016). The need to ensure food security, maintain soil productivity and protect agricultural products from damage and losses caused by plant pathogens, pests and weeds requires the extensive and continuous use of pesticides and inorganic/organic fertilizers, which in the long term have a detrimental effect on soil health, productivity and biodiversity (Prashar P., Shah S., 2016; Alengebawy A. et al., 2021). Soil microorganisms play a critical role in the functioning of terrestrial and subsurface ecosystems. They are involved in the formation of soil structure, carbon accumulation, decomposition of organic matter, nutrient cycling and, as a consequence, determine soil fertility and plant growth (Fierer N., Wood S.A., Bueno de Mesquita C.P., 2021). Therefore, the stability of soil microbial communities is one of the key factors that ensure the sustainable functioning of the entire soil ecosystem (Ramirez K.S. et al., 2018).

Ensuring the safety of both the environment and consumers necessitates a continuous search for innovative, effective crop protection products that are harmless to non-target organisms. In this context, environmentally friendly biological control of plant diseases has received considerable attention in recent years as an alternative to traditional chemical pesticides (Azizbekyan R.R., 2013). Biopesticides, which have proven to be highly effective in pest control and offer numerous advantages, are currently prominent (Chen J., 2018; Borges S. et al., 2021; Rajni Y., Singh S., Singh A.N., 2022). These natural substances, derived from plants and micro-organisms, provide a sustainable means of controlling phytopathogens (Kumar J. et al., 2021).

*Bacillus* bacteria are one of the major groups of soil and rhizosphere microbial pathogenic and opportunistic microbiota. *Bacillus* species have unique properties that make them promising candidates for the development of biological crop protection products. These properties include the ability to synthesize broad-spectrum antibiotics and antimicrobial compounds. It is this ability that makes them attractive as biological agents (Caulier S. et al., 2019; Baharudin M.M.A. et al., 2021; Xu B-H. et al., 2018). Bacteria of the *Bacillus* genus can be divided into two phylogenetically unrelated groups. One of these groups, the cereus group, includes human pathogens such as *B. anthracis* and food-borne pathogens such as *B.cereus*. These species are soil-dwelling and only *B. thuringiensis* is used in plant protection. Another group, “subtilis group”, including plant-associated species, such as *B. amyloliquifaciens, B. lichenoformis, B. pumilus,* and *B. subtilus* (Balleux G. et al., 2024). *B. velezensis* is a plant-associated species most of them registered for market as *B. amyloliquifaciens* (Dunlap C.A., 2019). *B. venezensis* genome contains gene clusters that encode for a wide variety of biosynthetic metabolites, including volatiles, syderophores, cyclic lipopeptides, and polyketides (Balleux G. et al., 2024).

Among the polyketide antibiotics, there are macrolactins. These are a relatively recently discovered group of macrolide antibiotics that have the genetic potential to be produced exclusively by *B.velezensis* (Steinke K. et al., 2021). Macrolactins are a class of 24-membered ring macrolides that are typically biosynthesized by a type I polyketide synthase. To date, more than 33 different macrolactins have been identified (Wu T., Xiao F., Li W., 2021).

With the development of biocontrol of plant diseases, macrolactins have also been found to be potent antimicrobial compounds that antagonize many bacterial plant pathogens and play an important role in suppressing plant diseases caused by *Agrobacterium tumefaciens* (Chen L., Xang X., Liu Y., 2021) *Fusarium oxysporum* and *Rhizoctonia solani (*) (Jaruchoktaweechai et al., 2000; Romero-Tabarez et al., 2006; Chen et al., 2009; Pandey et al. 2023).

The use of biologics derived from antibiotic-producing bacteria must be subject to strict risk assessment procedures as in the case of chemical formulations. However, resistome assessment and changes in the taxonomic composition of the microbiome are not currently used as part of the decision-making process for registration. In this case, it would be beneficial to conduct classical scientific research to provide an answer regarding the environmental consequences of using the biopreparation.

For example, the biological properties of cyclic lipopeptides have been well studied at various levels of the microbiome, from the molecular mechanisms of their antimicrobial action (Sur S. et al., 2018; Chen X. et al., 2022) to the taxonomic shifts in the composition of soil microbial communities (Kiesewalter H.T. et al., 2020; Yuan J. et al., 2017; Vasilchenko A.S. et al., 2025a). Much less is known about what happens to microorganisms at the cellular and conducted under controlled conditions that examines the effects of macrolactin A on soil microbiomes using 16S rRNA amplicon sequencing (Yuan J. et al., 2016). To gain a complete understanding of the impact of antibiotics on the indigenous microbiota, and consequently on soil health, a more in-depth study is required. This should include not only an assessment of taxonomic composition but also an analysis of the functional traits of the developing microbial community. In addition, there is a hypothesis that suggests that antibiotic production and antibiotic resistance are interdependent phenomena, designed to optimize the function of the microbiome by selecting beneficial partners (Spagnolo F. et al., 2021). Thus, profiling of antibiotic resistance genes (ARGs) in combination with the analysis of metagenome-assembled genomes allows us to obtain a list of species that are indicative of the microbial community when exposed to McA.

In our previous study, we isolated McA from a plant growth-promoting bacterium *Bacillus velezensis* strain X-Bio-1 and investigated the mechanism and spectrum of its antibacterial activity, including its ability to fight the phytopathogen *Pectobacterium carotovorum* (Vasilchenko A.S. et al., 2025b). To investigate the effects of McA exposure on the diversity, composition, stability, and function of soil microbial communities, and on the distribution of ARGs, DNA was extracted from soils treated with high and low doses of McA for 56 days, and from untreated controls. This study’s distinguishing feature is its use of real agrocenosis conditions. This assessment of the contribution and potential ecological risks of McA may inform the development of biopesticides based on this compound.

## 2. Materials and methods

### 2.1. Purification of McA

The *Bacillus velezensis* X-Bio-1 strain (Kravchenko S.V. et al., 2020) was cultivated in a 1.0-liter conical flask in a medium similar to LB broth (per liter of medium, g: tryptone −10, yeast extract −5, NaCl −2) for 48 hours. Bacterial cells were separated from the culture liquid by centrifugation and filtration through 0.22 µm membranes (VWR, US). Next, the antimicrobial component of the metabolites was extracted from the culture medium with ethyl acetate. The resulting extract was evaporated and re-dissolved in dimethyl sulfoxide (DMSO). Next stage of McA preparative purification was using low-pressure liquid chromatography system Buchi Reveleris X2-UV equipped by C18 cartridge (4.6 × 250 mm, 130 Å, 5 μm, Phenomenex, US). Quality of purified preparation was checked using reverse-phase HPLC on a Luna C18 analytical column (4.6 × 250 mm, 130 Å, 5 µm) (Phenomenex, US). The McA purity was >80%.

### 2.2. Experimental design

The studies were carried out on grey soils (Luvic Phaeozems, according to the WRB classification) Loam granulometric composition (clay 28.5%, silt 27.7%, sand 43.8%); рНН_2_О 6.2, рHКCl 5.6. The coordinates of the location where the soil samples were collected and the field experiment took place are: 57.0914, 65.3740. In the laboratory, the soil was homogenized, cleaned of visible plant debris and then sieved through a 2 mm sieve.

The experiment was conducted in a microcosm format, where the soil was prepared according to a specific scenario:1) control samples with the addition of sterile water (further referred as “control”); 2) experimental samples with the addition of McA solution at a concentration of 1 mg per kg of soil (further referred as “low dose”); and 3) experimental samples with the addition of sterile McA solution at a concentration of 10 per kg of soil (further referred as “high dose”). The prepared soil was transferred into polypropylene containers with multiple holes measuring 1–2 millimeters in diameter. These containers were filled with soil and placed at a depth of 0–5 centimeters in the agroecosystem (where the soil had been initially collected).

Each experimental group was represented in five biological replicates (microcosms), resulting in 15 metagenomic samples for the current analysis. The antibiotic-treated soil samples were collected 56 days after treatment. Soil samples were stored at −80^0^C until downstream molecular analysis.

### 2.3. DNA extraction and metagenomic sequencing

Total genomic DNA was isolated from 250 mg soil per sample using a Quick-DNA Fecal/Soil Microbe Kit (ZymoResearch, USA). DNA concentration and purity were determined using a Qubit 4.0 fluorometer (Thermo Fisher Scientific, USA) and a NanoPhotometer N120 spectrophotometer (Implen, Germany), respectively. Sequenced libraries were generated using the Illumina DNA Prep (M) Tagmentation Library Preparation Kit (Illumina, Inc., San Diego, CA, USA) according to the manufacturer’s instructions. Shotgun metagenomic sequencing was performed on the Illumina NovaSeq6000 platform (Illumina, Inc., San Diego, CA, USA) with a paired-end 2×150 read length protocol.

Low quality metagenomic reads were trimmed using Trimmomatic v.0.39 with paired default settings (Bolger A.M., Lohse M., Usadel B., 2014). The high-quality reads were *de novo* assembled using MEGAHIT v1.0.6 (Li et al., 2015) with default parameters. Taxonomic affiliation for each metagenome sample was performed with Kaiju v. 1.5.0 (Menzel P., Ng K.L., Krogh A.,2016). To identify metagenomic reads corresponding to antibiotic resistance genes, metagenomic reads were mapped to the Comprehensive Antibiotic Resistance Database (CARD) than 30% coverage were filtered out. ARG types/subtypes relative abundance was exhibited as ‘copy of ARG per copy of 16S rRNA gene’ (defined as ‘ratio’; Li et al., 2015). Assembled contigs were then annotated using the reCOGnizer v. 1.6.4 was used to identify clusters of orthologous groups (COGs) in the predicted protein sequences.

Genome binning from the metagenomic dataset was performed by Maxbin2 (Wu Y-W., Simmons B.A., Singer S.W., 2016), followed by dereplication and aggregation into metagenome-assembled genomes (MAGs) using DASTool (Sieber et al. 2018). The statistics of MAGs and the number of contigs were analyzed using QUAST (Gurevich A. et al., 2013). The completeness and contamination levels of MAGs were assessed using CheckM v. 1.1.3 (Parks et al., 2015), and only MAGs that met the criteria of >50% completeness and <5% contamination were selected for further analysis. The taxonomic affiliation and phylogenetic position of each MAG was determined using the GTDB-Tk (v2.4.0) tool and the Genome Taxonomy Database (GTDB) version r220. The phylogenetic tree was reconstructed using a *de novo* workflow with p Patescibacteria as the outgroup. The resulting phylogenetic tree was visualized and annotated using the Interactive Tree Of Life (iTOL) tool v7 (Letunic et al., 2021). Secondary metabolite biosynthetic gene clusters (BGCs) were identified in the MAGs using antiSMASH v5.2.0 (Blin K. et al., 2019).

### 2.4. Statistical analyses of dataset

Statistical analysis and visualization was performed in R version 4.4.1 environment. Most of graphs in this study were visualized using the “ggplot2” (Wickham H., 2016) package in R, with the exception of the co-occurrence network visualization which was created using Gephi.

The Alpha-diversity of bacterial genera were analyzed in terms of Shannon and Chao1 indexes using the microbiome v. 1.26.0 (Lahti et al., 2022) R packages. The beta-diversity analysis was conducted using the MicrobiotaProcess package in R (Xu et al., 2023). Permutational multivariate analysis of variance (PERMANOVA) on the Bray–Curtis dissimilarities was performed using the “mp_adonis” function. Principal coordinate analysis (PCoA) plots of Bray–Curtis dissimilarity or weighted UniFrac distances were created using the “mp_plot_ord” function in the MicrobiotaProcess (Xu, S. et al., 2023) package.

ARG profile similarities among samples were computed as Bray-Curtis distances and were visualized by PCoA. Adonis test was conducted to estimate the contribution of temporal, disinfection treatment and sample types on ARGs discrimination among samples using ‘mp_adonis’ function of MicrobiotaProcess package as described above.

Linear discriminant analysis Effect Size analysis (LEfSe) (Segata N. et al.,2011) was options (“multigrp_strat”), as implemented in the microbiomeMarker v. 1.6.0 (Cao Y. et al., 2022) package. Statistical significance was assessed using the Kruskal-Wallis test (p = 0.05) and Wilcoxon test (p = 0.05), both corrected for multiple comparisons using the False Discovery Rate (FDR).

Сo-occurrence network analysis of bacterial genera and ARG was calculated using integrated Network Analysis Pipeline (iNAP) (<u>Feng et al., 2022</u>). SparCC method was used to calculate the network with the following parameters: correlation strength exclusion threshold = 0.1, 100 shuffled times, threshold value > 0.7 and *p*-value = 0.05. Networks were visualized using Gephi 0.10.1 (Bastian M., Heymann S., Jacomy M., 2009).

## 3. RESULTS

### 3.1. The Impact of McA on soil microbial community, diversity, composition, and stability

Shotgun metagenomic sequencing of 15 soil samples was performed using the Illumina NovaSeq platform 6000. DNA from untreated control soil and soil exposed to a low or high dose of McA was used to generate three sequence data sets. A total of approximately 1.43×10^9^ reads were generated, with an average of 9.5×10^7^ reads per sample for the soil experiments (Table S1).

The alpha diversity of soil bacterial taxa, measured by the Shannon diversity index and by Chao1 richness, was not significantly affected by McA exposure (Fig. 1A). A comparison of microbial community beta-diversity using the Bray-Curtis dissimilarity index revealed the effect of soil treatment with McA on clustering ordination. Both McA concentrations, low and high, had a significant effect on the diversity of the soil microbial community after 56 days of exposure - the soil sample clusters were closer together (highlighted by the green and yellow ellipses) than the control biome sample cluster (Fig. 1B).

**Figure 1.**
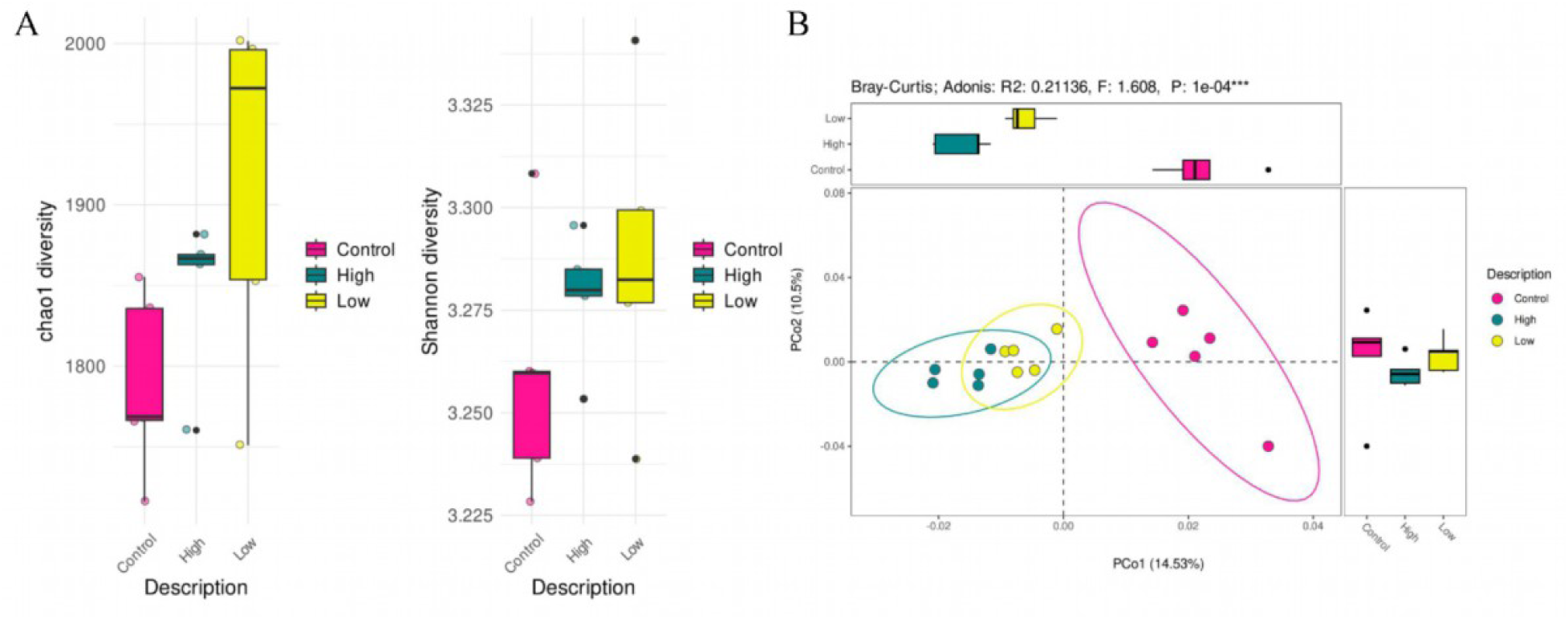
The diversity of bacterial communities in soil under different McA treatments. **A)** Alpha diversity (Shannon index) and richness (Chao1). **B)** Principal component analysis (PCoA) visualization of the pairwise dissimilarity (Bray-Curtis distance) in the bacterial communities.

We characterized changes in the diversity and compositions of the bacterial community across the McA treatments. At the phylum level, we identified 44 bacterial taxa across the 15 soil samples (all detected phyla are listed in Table S2). Actinomycetota, Pseudomonadota, Bacillota, Planctomycetota, and Myxococcota were the dominant taxa in soils treated with different doses of McA and untreated soils. Analysis of the relative abundance of microbial phyla showed that Actinomycetota abundance was reduced by both high (64.23%) and low (64.83%) doses of McA compared to control samples (65.97%) (p < 0.05). The relative abundance of the other prevalent taxa, *i.e*., Planctomycetota and Myxococcota, also decreased significantly in response to both high and low doses of McA (p < 0.05). Conversely, the relative abundance of Pseudomonadota increased with both high (31.88%) and low (31.04%) doses of McA compared to untreated samples

The relative abundance of Bacillota was similar across all experimental groups (2.57%, 2.34%, and 2.68% for low, high, and control groups, respectively; Fig. 2A).

**Figure 2.**
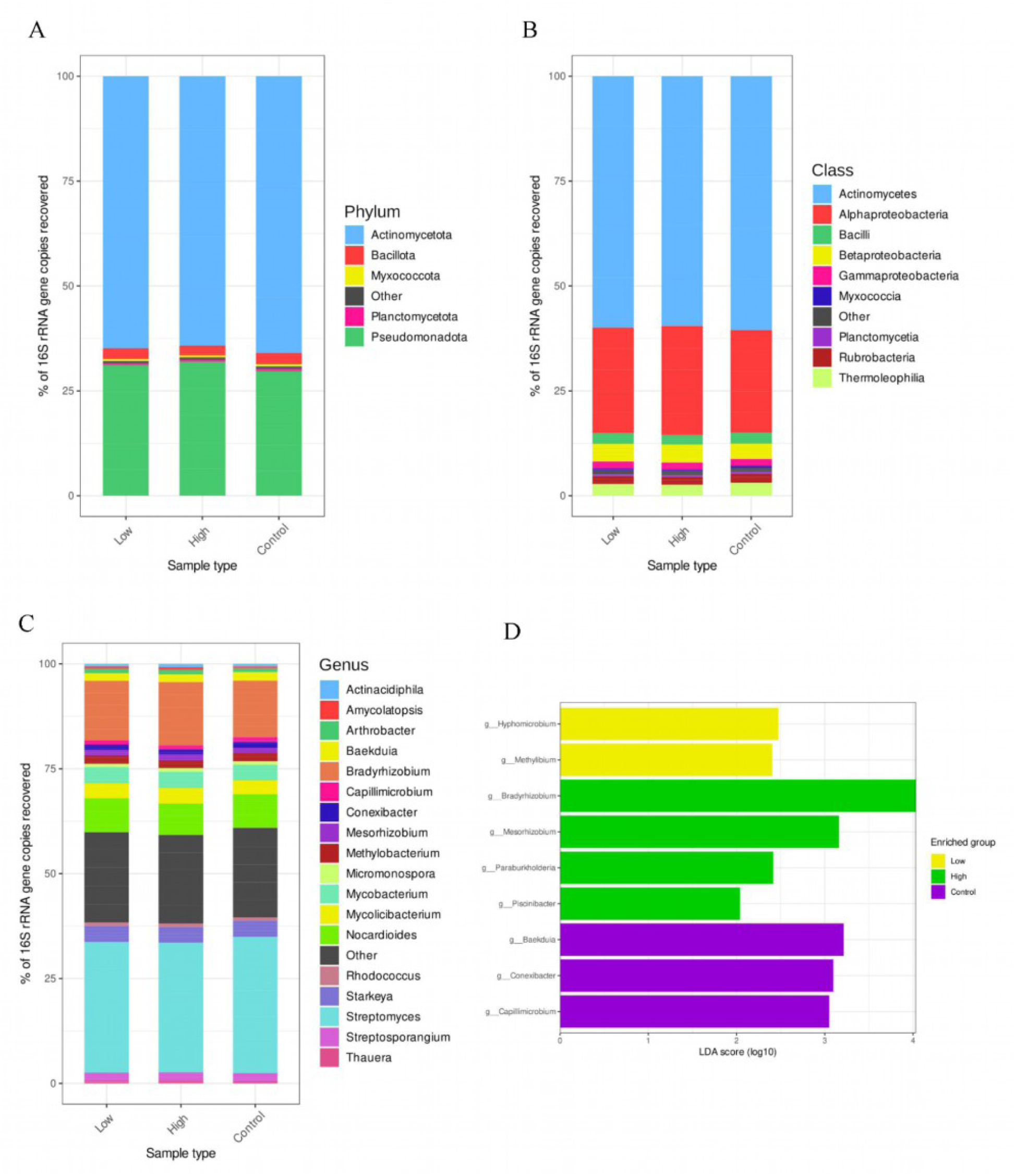
Composition of bacterial communities at phylum **(A)**, class **(B)**, genus **(C)** level in soil samples from different McA treatments. LEfSe analysis reveals changes in the bacterial genera between the three experimental groups **(D).**

Taxonomic profiling of all samples revealed 103 bacterial classes, with Actinomycetes, Alphaproteobacteria, Betaproteobacteria, Gammaproteobacteria, Thermoleophilia, Bacilli and Rubrobacteria being dominant and representing >98% of the total bacterial fraction (Fig. 2B). Exposure to high and low doses of McA significantly decreased the relative abundance of several dominant taxa compared to the control group: Thermoleophilia (from 3.14% to 2.63% and 2.79%, respectively; p < 0.05) and Rubrobacteria (from 2.09% to 1.85% and 1.89%, respectively; p < 0.05). Among the minor taxa, significant decreases were observed in Planctomycetia (from 0.50% to 0.44% and 0.45%; p < 0.05) and Acidimicrobia (from 0.17% in the control to 0.16%; p < 0.05) after exposure to both high and low doses of McA. In contrast, McA significantly increased the relative abundance of Alphaproteobacteria (from 24.38% to 25.92% and 25.13%; p < 0.05) and Betaproteobacteria (from 3.70% to 4.33% and 4.24%; p < 0.05) at both high and low doses. In addition, high McA concentration significantly reduced the relative abundance of Actinomycetes (60.52% vs. 59.56%; p < 0.05) and Myxococci (0.53% vs. 0.44%; p < 0.05).

The thirteen dominant bacterial genera (each comprising >1% of the total community) across all three groups were *Streptomyces, Bradyrhizobium, Nocardioides, Starkeya, Mycobacterium, Mycolicibacterium, Methylobacterium, Streptosporangium, Baekduia, Mesorhizobium, Conexibacter, Arthrobacter, and Capillimicrobium* (Fig. 2C). These genera comprised approximately 75% of the total soil bacterial community. There were significant differences (p < 0.05) in the composition of the soil microbial communities after the McA treatment (Fig. 2D). The analysis of the significant effects of McA on the relative abundance of microbial genera showed: a decrease of *Baekduia* (from 2.08% to 1.83% and 1.87%), *Conexibacter* (from1.32% to 1.13% and 1.20%), and *Capillimicrobium* (from 1.15% to 0.98% and 1.02%) after the high and low McA treatment. In addition, we observed a reduction in abundance of *Paraconexibacter* (from 0.52% to 0.38% and 0.43%) and *Paludisphaera* (from 0.09% to 0.07% and 0.08%) after high-dose McA treatment (p<0.05). After McA treatment, the genera that significantly increased (p<0.05) in abundance differed between the high and low dose treatments. High doses of McA increased the abundance of *Bradyrhizobium* (from 13.46% to 15.06%), *Mesorhizobium* (from 1.26% to 1.47%), *Paraburkholderia* (0.23% vs. 0.27%), and *Piscinibacter* (0.04% vs. 0.05%), while *Hyphomicrobium (*0,01% vs. 0,06%) and *Methylibium (0,01%* vs. 0,05%) were more abundant at low McA doses than in controls.

A LefSe-based analysis identifies bacterial taxa that can be considered potential biomarkers for certain conditions, particularly the structure of the soil microbiome affected by McA. These taxa were identified as having an LDA score greater than 2, indicating that they can be used as indicators of changes under McA action. The four genera identified as indicators for high McA dose were *Bradyrhizobium, Mesorhizobium, Paraburkholderia,* and *Piscinibacter.* Two genera were found to be indicators of microbial composition under low doses of McA, and three genera represent the taxa of the normal state composition (Fig. 2 D).

### 3.2. Antibiotic resistance genes in soil metagenomes

This study was able to compare the relative abundance of ARGs on McA-treated and untreated soils in the microcosms. To investigate differences in soil ARG characteristics under varying McA concentrations, we performed principal coordinate analysis (PCoA) on ARGs abundance data using the Bray-Curtis dissimilarity index (Fig. 3A). PCoA revealed that McA treatment explains 22.6% of the changes in the distribution of soil ARGs (PERMANOVA, R² = 0.226, p = 0.05). Treatment of soil samples with high and low doses of McA did not result in statistically significant differences in ARG richness (Chao1 index). Similarly, no effect was found for the ARG’s Shannon diversity index (Fig.3B).

**Figure 3.**
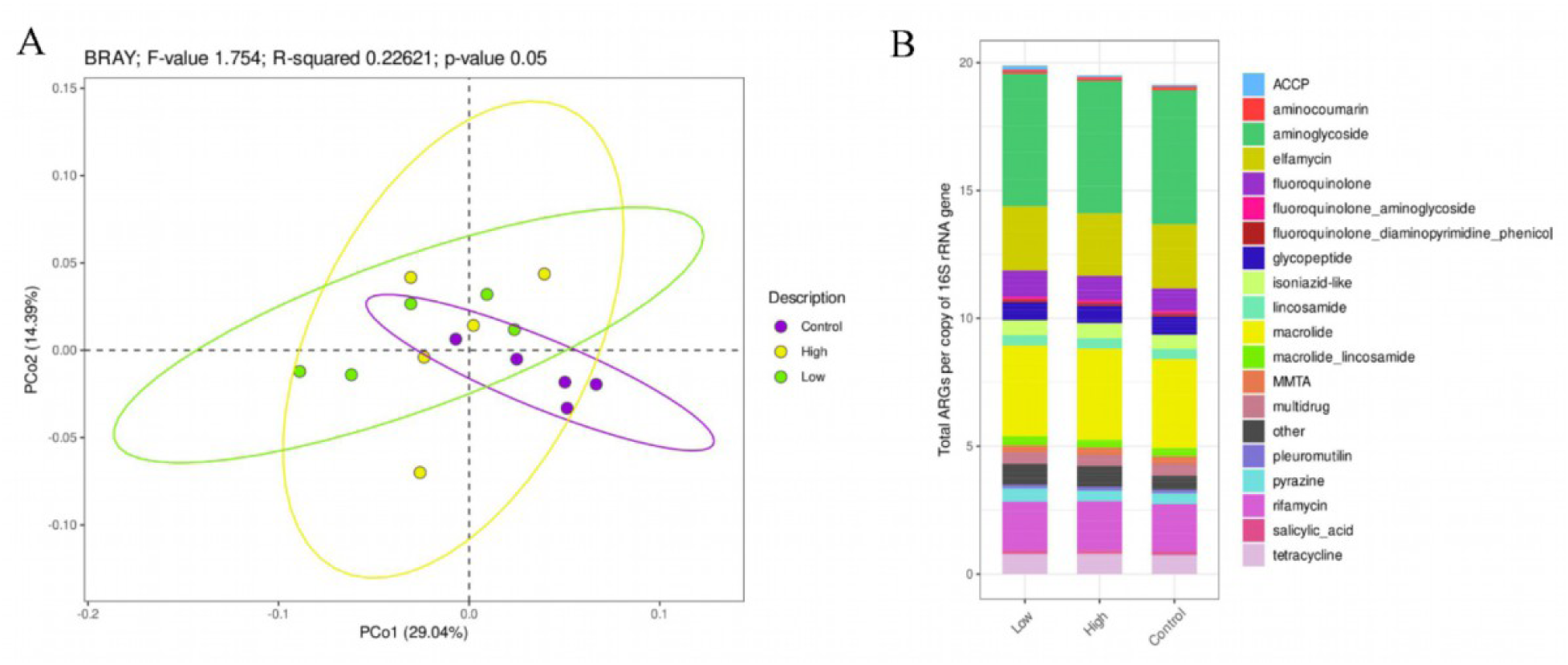
The diversity of soil resistomes under different McA treatments. **A)** Beta diversity was estimated using the Bray-Curtis distance-based PCoA. **B)** Alpha diversity was estimated using Chao1 and Shannon indices diversity of the composition of antibiotic resistance genes in soil.

Profiling the ARG composition it was found that soil resistome comprised 49 antibiotic classes. Aminoglycosides, macrolides, elphamycin, rifamycin and fluoroquinolones were the most common classes of antimicrobials, accounting for more than 50% of the total ARGs detected (Fig. abundant ARGs and accounted for more than 1% of the total number of ARGs (Fig. S 3). Genes conferring resistance to aminoglycosides (12%), macrolides (15%), elfamycins (6%) and rifamycin (11%).

Analysis of the differential gene representation in the comparison groups showed that soils treated with a low dose of McA had significantly higher levels of genes (n=7) responsible for resistance to glycopeptides (*vanHO*), fluoroquinolones (*mfpA*), elphamycins (*facT*, conferring resistance to factomycin), macrolide_ fluoroquinolones (*mexY*), isoniazid-like antibiotics (*Mtub_inhA_INH*), macrolides_lincosamides_streptogramin (*Erm(O)-lrm*) and phenicol antibiotics *(cmlv*) (Fig.4B).

**Figure 4.**
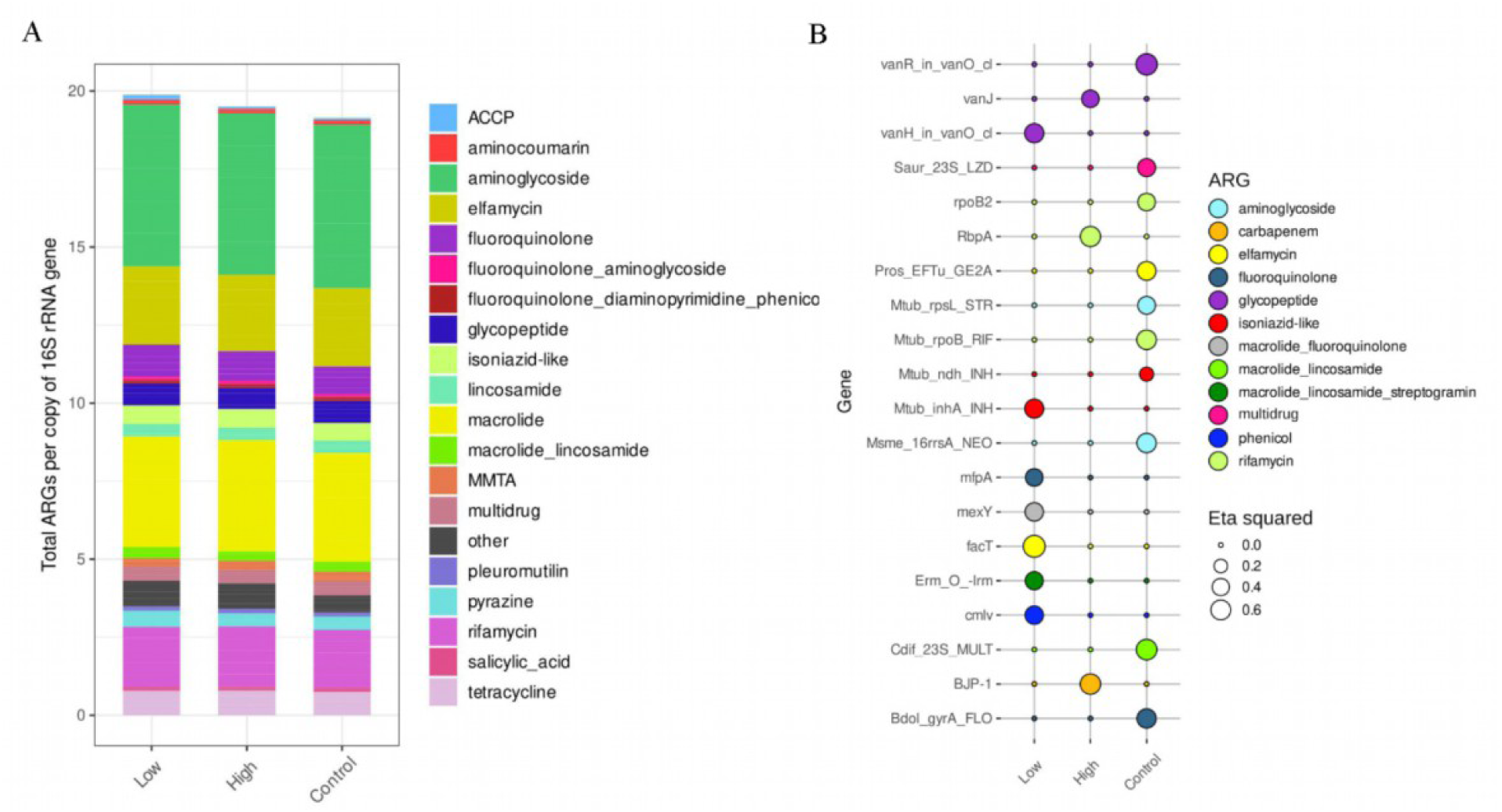
The composition and differences of ARGs in soil under different treatments of McA. **A)** The compositions of ARG type in different experimental groups. **B)** Differentially expressed ARGs in response to McA exposures according to LefSe analysis.

Only three genes conferring resistance to carbapenems (*BJP-1*, which hydrolyzes most beta-lactams), rifamycins (*RbpA*, which confers resistance to rifampin), and glycopeptides (*vanJ*, which confers resistance to teicoplanin) were found to be enriched in the soil treated with high doses of McA (Fig. 4B). In the control soil, the relative abundance of ten antibiotic resistance genes increased significantly (p<0.05). These included genes conferring resistance to aminoglycosides (*Msme_16rrsA_NEO*, neomycin; *Mtub_rpsL_STR*, streptomycin); rifamycin (*rpoB2*, rifampin; *Mtub_rpoB_RIF*, rifampicin); elfamycin (*Pros_EFTu_GE2A*); glycopeptides (*vanR_in_vanO_cl*); macrolide-lincosamides (*Cdif_23S_MULT*); multidrug (*Saur_23S_LZD*, linezolid); isoniazid (*Mtub_ndh_INH*); and fluoroquinolones (*Bdol_gyrA_FLO*) (Fig.4B)

To investigate potential host microorganisms of ARGs, this study used correlation analysis between the abundance of each genus and ARGs in the community, assuming that genera with a high correlation with ARGs are potential host microorganisms. To identify relationships between microbes and antibiotic resistance genes (ARGs), we applied correlation analysis and visualized the data using Gephi, displaying patterns of co-detection of ARGs and microbial communities (Fig. 5). The co-occurrence network was comprised of 86 nodes (52 ARGs and 34 bacterial genera) that formed 185 links. Among these links, 88.64% were positively correlated, and 11.36% were negative (Fig. 5).

**Figure 5.**
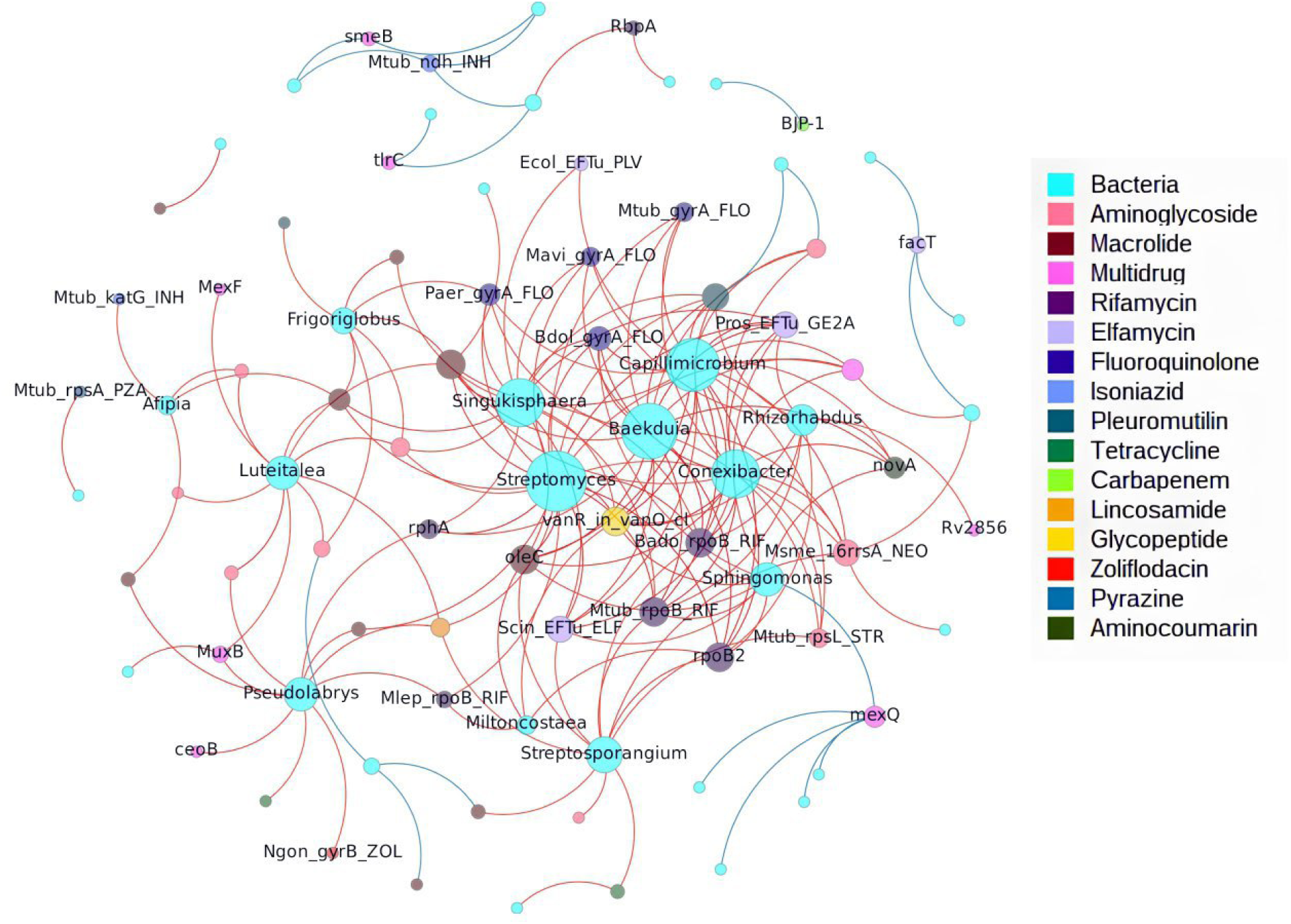
Co-occurrence network showing the association of affected ARGs with bacterial genera (Spearman’s R >0.7 and p *<* 0.01). Node size is proportional to the number of edges. Red and blue links indicate positive and negative significant correlations between the abundance of ARGs and the prevalence of bacterial genera in the samples. In the network, bacteria are represented by the blue nodes, ARGs by the colored by the class of antibiotic.

A total of 22 bacterial genera had significant positive correlations with ARGs, among which *Streptomyces*, *Baekduia*, *Capillimicrobium*, *Conexibacter* from *Actinomycetota*, and *Singulisphaera* from *Planctomycetota* carried more diverse ARGs (21, 19, 18, 16, and 16, respectively) than other one. For example, *Streptomyces, Baekduia, Capillimicrobium* and *Conexibacter* were found to have positive and significant connections with resistance genes against rifamycin (*rphA, rpoB2, Bado_rpoB_RIF, Mtub_rpoB_RIF*), elfamycin (*Scin_EFTu_ELF*), fluoroquinolone (*Mtub_gyrA_FLO, Paer_gyrA_FLO, Bdol_gyrA_FLO, Mavi_gyrA_FLO*), macrolide (*Ctra_23S_MAC*), glycopeptide (*vanRO*), and neomycin (*Msme_16rrsA_NEO*). Notably, the dominant taxa *Nocardioides* and *Bradyrhizobium* were found to be potential carriers of only one resistance gene, *rpsA* and *rbpA*, respectively. The details of co-occurrence pattern between ARG subtypes and bacterial taxa at genus level were presented in Table S5.

### 3.3. Effect of McA on functional traits of bacterial community of soil

The most abundant functional categories (COGs) were amino acid transport and metabolism, energy production and conversion, lipid transport and metabolism, replication, recombination and repair, translation, ribosomal structure and biogenesis. Lefse analysis results show COG terms that were differentially expressed between the treated samples in comparison to untreated samples. More than 20 COGs were enriched in the control group compared to McA-treated (Figure 6). These genes are primarily involved in replication and recombination, repair processes (COG0587, COG2176, COG0556), signal transduction (COG2114, COG2206, COG4585), lipid transport and metabolism (COG2409, COG0365, COG4770), and amino acid transport and metabolism (COG0160, COG1167). The control group showed an increase in ABC-type multidrug transport system (COG1131) proteins (Fig. 6). Exposure of soil to high doses of McA increased the abundance of the gene responsible for the biosynthesis, transport, and catabolism of secondary metabolites (COG3321), namely polyketide synthase (PKS) enzymes, inorganic ion transport and metabolism (COG0715), amino acid transport and metabolism (COG0520, COG0403). Exposure to low concentrations of McA increased the abundance of genes associate with signalling, chemotaxis (COG0784 and COG0840) and broad substrate specificity drug efflux (COG0841, COG0845, COG1538) (Figure 6).

**Figure 6.**
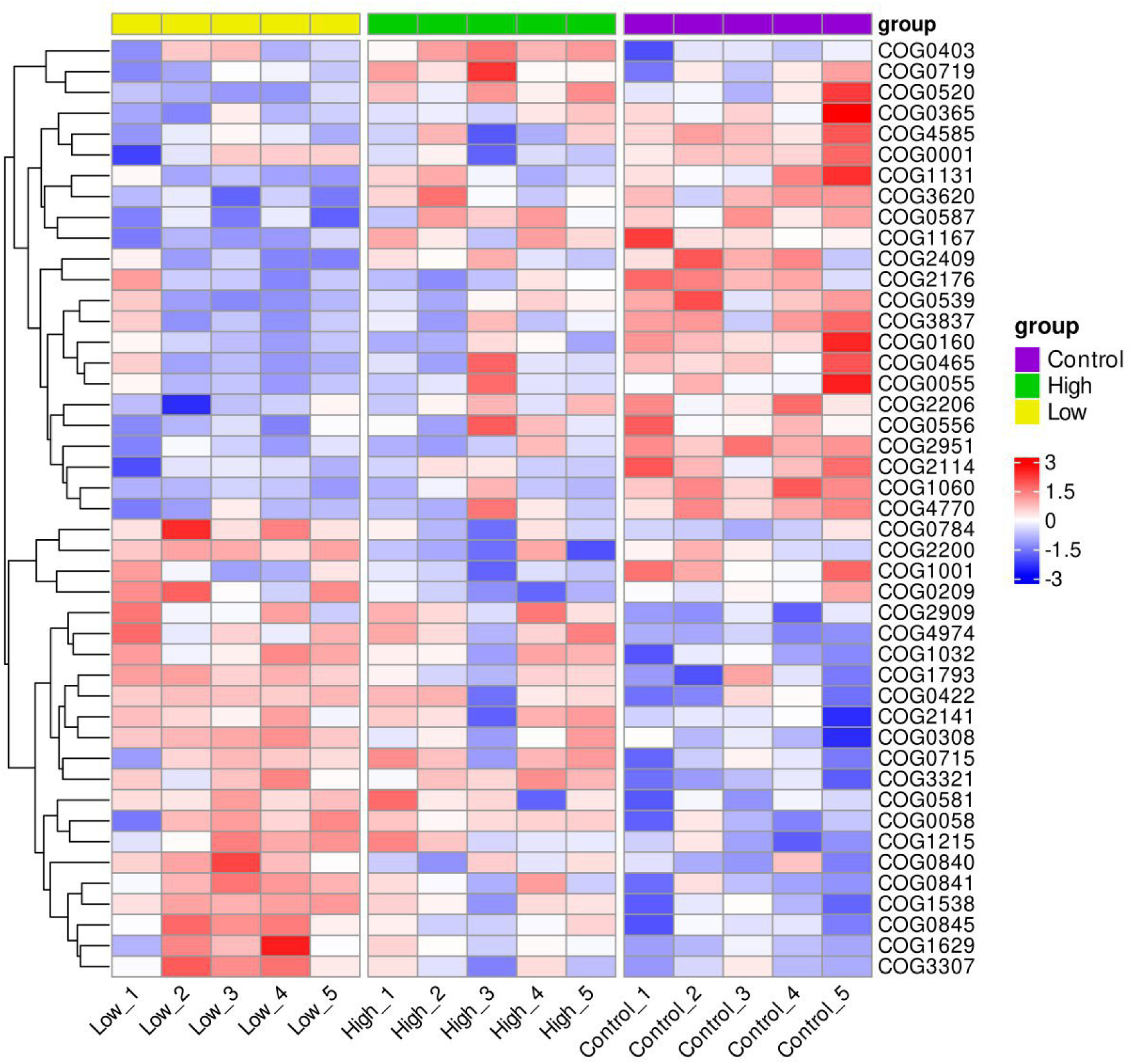
Changes in the potential functional activity of soil microbiomes under the influence of MсA. Lefse-analysis of the relative abundance of functional clusters of genes (COGs) involved in various biological processes in soil microcosms.

### 3.4. Metagenome assembled genomes of antibiotic-resistant bacteria

We reconstructed 91 metagenome-assembled genomes (MAGs) from the shotgun metagenomes generated in this study. All MAGs were classified into 3 categories: high quality assembly (completeness >90% and contamination <5%), medium quality (completeness >50% and <90%, contamination <10%) and low quality (contamination >5% and <25% and completeness >50%) according to the Minimum Information on Metagenome Assembled Genome (MIMAG) standard. Only one genome assembly was found to be of high quality, while 45 MAGs were classified as genomes of medium assembly quality and 45 MAGs as genomes of low the GTDB. Two phyla were identified, with the most abundant being Pseudomonadota, including class Alphaproteobacteria (n=48), Gammaproteobacteria (n=2) and Betaproteobacteria (n=1). The phylum Actinomycetota was represented only by the class Actinomycetes (n=30).

Thirty-seven metagenome-assembled genomes (MAGs) of high and medium assembly quality were used for further analysis. A phylogenetic tree was constructed for them (Fig. 7). Among the reconstructed MAGs, Alphaproteobacteria (n=16) were the most abundant and represented by the following genera: *Pseudolabrys*, *Methylovirgula*, *Hyphomicrobium*, *Methyloceanibacter*, and a member of the family Xanthobacteraceae not identified to genus (Table S6). Only two MAGs represented the class Gammaproteobacteria. Class Binatia (n=11) included the DP-1 genus. Actinomycetes (n=1) was represented by the Propionibacteriaceae sp. Two MAGs belonging to the genus JACDDX01 within the class Gemmatimonadetes. Class Thermoleophilia represented by the *Solirubrobacterales* spp.

**Figure 7.**
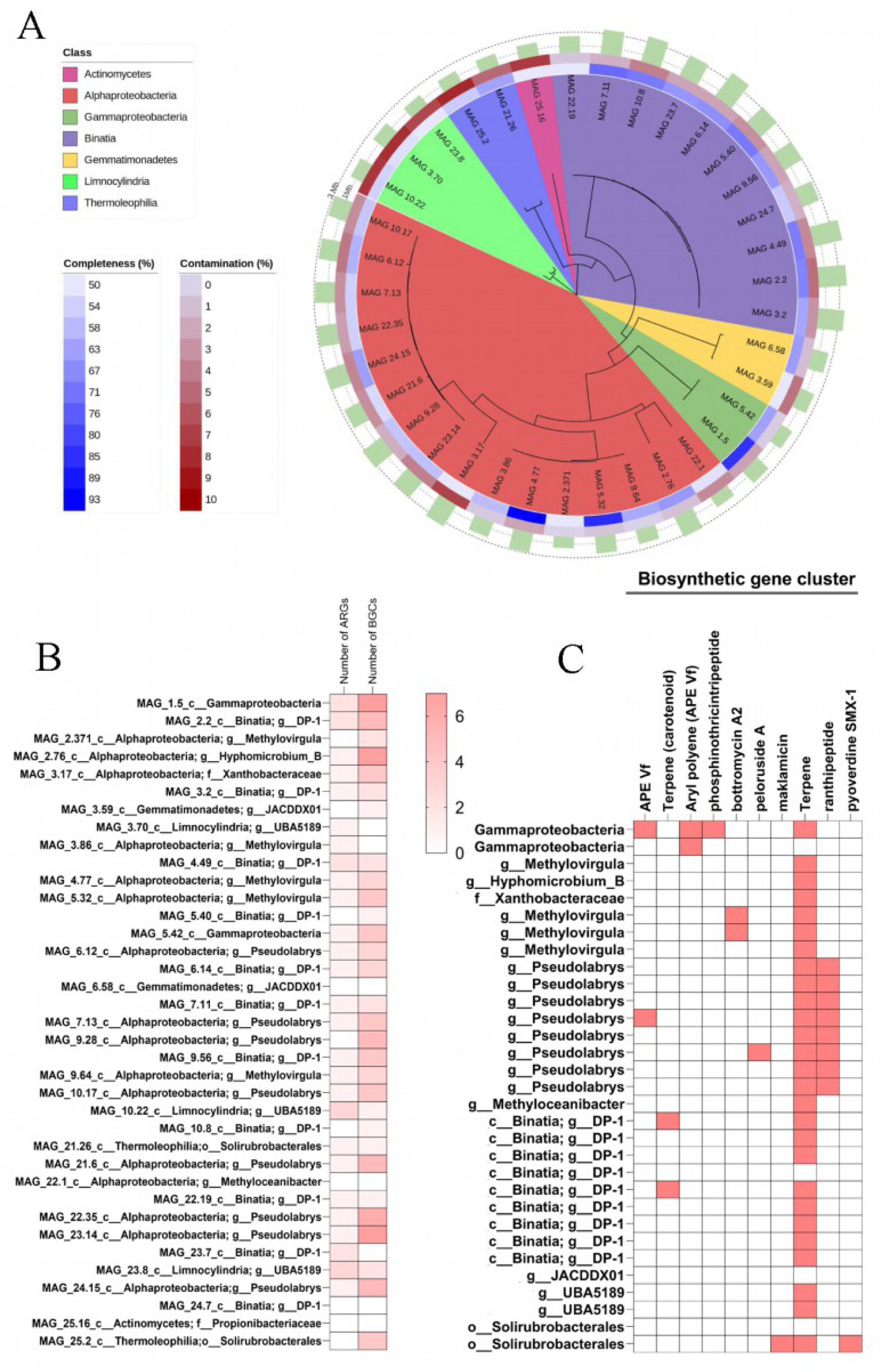
**A)** Phylogenetic visualization of the recovered MAGs of high and medium quality at class-level MAGs (n = 37); **B)** The number of ARGs is shown alongside the biosynthetic gene clusters (BGCs) involved in protection, siderophore production, and antibiotic biosynthesis.

These 37 metagenome-assembled genomes (MAGs) contained a total of 35 antibiotic resistance genes (ARGs) across 8 distinct subtypes (Table S7). More than half (54%) of the MAGs carried genes conferring resistance to tetracycline (*adeF*) and vancomycin (*vanT, vanY*). Additionally, ARGs identified in MAG_1.5_ Gammaproteobacteria spp. conferred resistance to benzalkonium chloride (*qacG*), trimethoprim and chloramphenicol (*rsmA*). The greatest diversity (4 different ARGs) of ARGs was found in members of the phylum Chloroflexota (class Limnocylindria), which carried resistance genes to glycopeptide (*vanYB, vanYG, vanWI*) and tetracycline (*otrA*) antibiotics. In ten of the 37 MAGs (Figure 7), no antibiotic resistance genes were detected (Fig.7).

We also analyzed the presence and diversity of putative biosynthetic gene clusters (BGCs) responsible for the biosynthesis of secondary metabolites. Among 31 MAGs, we identified 106 putative BGCs, 11 of which showed negligible similarity to known clusters. The vast majority of MAGs (31 out of 37) had at least one BGC. These BGCs encoded biosynthetic products predominantly belonging to the following classes: terpenes (27), RRE-element-containing clusters (15), ranthipeptides (8), redox-cofactor clusters (8), type III polyketide synthases (T3PKS, 6), non-ribosomal peptide synthetases (NRPS, 6), and others (Table S8). Alphaproteobacteria were the primary contributors to these BGC types (Table S9). These BGCs contained genes for the biosynthesis of various antibiotics or antimicrobial substances (terpenes, bottromycin A2, phosphinothricin tripeptide, maklamicin, peloruside A, ranthipeptides), siderophores (pyoverdine SMX-1), and protective substances (carotenoids, hopanes, aryl polyenes [APE Vf]) (Fig. 7B).

## 4. Discussion

The use of synthetic agrochemicals with antibiotic properties is currently the most effective approach to controlling phytopathogenic bacteria (Ali N. et al., 2016). Overuse of harmful agrochemicals, sometimes inappropriately, negatively impacts soil ecology and the wider environment (He et al., 2008). Large amounts of pesticides entering the soil have a direct impact on the soil microbiota, a biological indicator of soil fertility that affects plant growth and development (Onwona-Kwakye M. et al., 2020).

This scenario leads to a constant search for innovative, environmentally friendly products to protect crops while avoiding harm to non-target organisms and remaining safe for human consumption. In recent years, there has been a lot of focus on natural-derived substances and biological agents for pest control (Azizbekyan R.R., 2013). Broad-spectrum natural antimicrobials are preferred for pathogen suppression because they can suppress multiple pathogens simultaneously. However, this broad-spectrum activity can also have serious unwanted effects on the soil bacterial community, as many non-target bacteria, including beneficial bacteria, can be killed at the same time. Our previous study demonstrated that *Bacillus velezensis* X-Bio-1 effectively inhibited the growth of narrow spectrum of bacterial phytopathogenes. This antibacterial activity was attributed to the production of the secondary metabolite macrolactin A (Vasilchenko et al., 2025b).

In this study, we have found that the beta diversity of the soil bacterial community changes upon exposure to McA at low and high doses (Fig. 1B). After assessing the taxonomic changes, we conclude that McA has a significant impact on the bacterial community, even at the phylum level. Among these taxa, actinobacteria are reduced under the influence of antibiotics, while the representation of the Pseudomonadota phylum increases. Both taxa play an important role in microbiological processes, both in soil and in the rhizosphere of plants (Rutere et al. 2020; Vio S. A. et al., 2020; Zhang et al. 2023).

A similar significant decrease in the abundance of Acidobacteria, Actinobacteria, and a significant increase in Proteobacteria (Pseudomonadota) was observed in the soil bacterial community after one-month treatment with mixture containing three macrolactin compounds (macrolactin A, 7-O-malonyl macrolactin A, and 7-O-succinyl macrolactin A) (Yuan J. et al., 2016). Although Actinomycetota has been reported to have a high resistance to environmental stress (Gao B., et al., 2012), based on our results, we can conclude that they are sensitive to McA. At the genus level, among the 13 dominant bacterial genera (Table S4), the presence of 3 genera Gram-positive *Baekduia, Conexibacter*, *Capillimicrobium* was reduced when exposed to McA, which showed broad spectrum antibacterial activity of McA. It is important to note here that antibacterial activity in respect to the soil microbiome can be realized indirectly through changes in microbe-microbe relationships. Our experience in studying the effects of antimicrobial substances, in particular cyclic lipopeptides, indicates that substances that do not possess bactericidal or bacteriostatic properties can significantly impact the composition of bacterial communities in soil (Vasilchenko et al. 2025b).

On the other hand, the relative abundance of dominant bacterial genera belonging to Pseudomonadota (Alphaproteobacteria and Betaproteobacteria) was significantly increased by the addition of McA. In particular, the relative abundance of the major genera *Bradyrhizobium* and *Mesorhizobium* has been significantly increased, indicating that these soil bacteria benefit from McA addition (Table S4). *Bradyrhizobium* species are important member of soil community which perform a wide range of biochemical functions including nitrogen fixation during symbioses, denitrification and aromatic compound degradation (Okubo et al., 2012; Serbent et al., 2019; Huang et al., 2017). The multiple roles of *Bradyrhizobium* in the nitrogen cycle make the ecology of this group important for agriculture (Jones, F., Clark, I., King, R. *et al*., 2016). *Mesorhizobium*, plant growth-promoting rhizobacteria (PGPR), produce indole-3-acetic acid (IAA), ACC deaminase, and fix nitrogen (Verma et al., 2013; Lemaire et al., 2015b). Most of the genomes reconstructed from the metagenomes of McA-treated soils belong to the phylum Pseudomonadota, mainly to the classes Alphaproteobacteria and Gammaproteobacteria. Alphaproteobacteria, represented by the genera *Pseudolabrys* and *Methylovirgula*, contribute to soil nitrogen fixation (Li Z. et al., 2022) (Table S5).

An important aspect of studying the effects of antibiotics on natural habitats is the analysis of resistomes, which is a set of genes that encode antibiotic resistance (Wright G.D., 2007). A lot of research has been done on how the use of antibiotics affects the soil resistome. Most such studies focus on industrial wastewater, which contains medical-grade antibiotics (Lai F.Y. et al.,2021; Zhao Y. et al.,2020; Delgado-Baquerizo M. et al.,2022). Our study focuses on an antibiotic McA that is part of the metabolome of soil and rhizosphere bacilli. The use of metagenomic sequencing has allowed us to not only identify qualitative and quantitative changes in the composition of ARGs, but also to determine the hosts of these genes.

The low dose of McA altered the numerical diversity of the resistome as the number of unique antibiotic resistance genes increased (Fig. 4C). According to recent data, McA has a mode of action similar to that of elfamycin, that is, it alters the elongation factor EF-Tu in bacteria (Vasilchenko et al., 2025). In this regard we have paid attention to the abundance of ARGs which conferring resistance to elfamycins. We identified across all groups 3 ARGs which determined found that control soils have more copies of the *Pros_EFTu_GE2A* gene, while McA-treated soils have more copies of the *facT* gene. Thus, the data suggest that exposure to low concentrations of McA selects for bacteria whose genome encodes proteins that determine resistance to the elfamycin antibiotic, factumycin (Thaker M.N. et al. 2012).

We used two methods to find hosts for different ARGs. The first method is correlation analysis with a high threshold level (Rs>0.7). The second method is the annotation of high- and medium-quality MAGs. According to correlation analysis, the hosts for elfamycin-related ARGs were identified as *Baekduia, Capillimicrobium, Conexibacter, Singulisphaera, Sphingomonas*, and *Streptomyces*. Among these, no taxa were found that are prevalent in McA-treated soils (see Table S5). Among MAGs, we also did not find any of these species or elfamycin-related ARGs. Analysis of others ARGs in MAGs also revealed no genetic determinants of resistance to macrolides. Thus, we were unable to establish a correlation between the enrichment of soil with specific taxa under the influence of McA and the presence of elafamycin resistance genes in the resistome. Instead, we found that the number of genes resistant to GE2270A decreased and the number of genes resistant to factumycin increased. Due to the lack of detailed information on the differences in the mechanisms of action between McA and other elfamycins, it is still difficult to adequately integrate the data obtained from this study.

It is surprising that only three antibiotic resistance genes (*RbpA, vanJ*, and *BJP-1*) were found to be enriched in microcosms with a higher McA dose. As for other ARGs, the following ARGs were found during annotation of MAGs: the *adeF* gene encoding the membrane fusion protein AdeFGH, which is part of the multidrug efflux complex and confers resistance to tetracycline and fluoroquinolones, the *rsmA* gene conferring resistance to the diaminopyrimidine classes, phenicols and fluoroquinolones, the *qacG* gene encoding a small multidrug resistant efflux pump conferring resistance to benzalkonium chloride and ethidium bromide (Rui Y.; Qiu, G., 2024).

In general, variations in ARGs were considered to be closely related to shifts in bacterial community composition, a potential driving factor for the emergence of the resistome (Zhao R. et al.,2018). Our correlation analysis revealed a significant positive correlation between the bacterial communities and ARGs. In this study, Actinomycetota (*Streptomyces, Baekduia, Capillimicrobium, Conexibacter*), Pseudomonadota (*Rhizorhabdus, Pseudolabrys, Sphingomonas*), and *Singulisphaera* from Planctomycetota were the prevailing potential ARG hosts (Fig. 5). Notably, the dominant taxa *Nocardioides*, *Bradyrhizobium* and *Mesorhizobium* were found to be potential carriers of only one resistance gene, *rpsA* and *rbpA*, respectively. Some potential ARG hosts have been verified in previous studies. The self-resistance in antibiotic-prevailing and threatening in various environments at the present time (Ogawara H., 2016). *Planctomycetota* have also been detected by metagenomic approaches in hotspots for AMR dissemination such as wastewater treatment plants (Cayrou C. et al., 2013; Aghnatios R., Drancourt M., 2015).

According to the hypothesis, as proposed by Spagnolo, Trujillo and Dennehy (2021), the synthesis of antibiotics and resistance to them forms a paired relationship that affects the assembly of a microbial community. In this context, we set out to evaluate how the functional potential of the soil microbiome changes when exposed to McA.

According to the COG annotation and clustering results, some COG terms have different enrichment patterns in McA treated and untreated soils (Fig. 6). For example, COG3321, an enzyme involved in polyketide biosynthesis (PksD), was enriched in the high-dose McA group. In addition, high dose of McA promoted the enrichment of the other COG terms describing transport and metabolism processes in bacteria, including inorganic ions (COG0715), amino acids (COG0520, COG0403) and carbohydrates (COG0058). The low dose of McA enriched multidrug-efflux transport proteins (COG0841, COG0845), response regulator receiver protein (COG0784). The relative abundance of cell wall/membrane/envelope biogenesis genes/proteins (COG1538, COG3307) in the low-dose McA treatment was significantly higher than in the untreated control. Bacteria use various strategies to adapt to complex environmental conditions, with BGCs playing a crucial role in the production of various compounds that not only facilitate their environmental adaptation and give them a competitive advantage, but also contribute broadly to the evolutionary pressures and ecological dynamics that shape ecosystems (Dong X, et al., 2024).

## 5. Conclusion

Thus, the study found that McA, when introduced into the soil, had a significant effect on the bacterial community. However, from the perspective of changing the composition and abundance of antibiotic resistance genes, McA did not result in an increase in the number or diversity of ARGs Against the background of an increase in the proportion of some genes, the number of other genes decreased, so the overall diversity of genes remained the same. The intriguing looks like an increase in the dose of McA by ten times did not lead to a proportional increase in the effects. An analysis of the representation of functional genes also suggests that the dose of 1 mg per kg of soil has a more significant effect than a higher dose. All this data needs additional research on soils with different physical and chemical properties in order to understand how the effects of McA can be different in relation to different soil habitats.

## CRediT authorship contribution statement

**Darya V. Poshvina -** Writing – original draft, Data curation, Conceptualization, Formal analysis, Funding acquisition. **Alexander S. Balkin** – Investigation, Software, Visualization, Formal analysis. **Diana S. Dilbaryan** – Investigation, Formal analysis. **Alexey S. Vasilchenko** - Writing – review & editing, Conceptualization, Validation.

## Data availability

The raw shotgun sequencing datasets and the sequence data for 37 MAGs have been deposited into the NCBI Sequence Read Archive (SRA) in the National Center for Biotechnology Information (NCBI) database under the Bioproject PRJNA1107916.

## Acknowledgments

This work was supported by the Ministry of Science and Higher Education of the Russian Federation for financial support (FEWZ-2024-0005) in parts devoted to McA purification and annotation of microbiome functional properties: sections 3.3, 3.5.

