## Supplemental Data 1 for "Responses of Soil Bacterial Community and Its Resistome to Short-term Exposure to Macrolide Antibiotic Macrolactin A: Metagenomic analysis": Supplementary files.docx

**Table S1.** Summary of metagenomic sequencing data for different sample groups

|  | Number of reads (Gb) | Number of contigs | Largest contig | Total length(bp) | N50 |
| --- | --- | --- | --- | --- | --- |
| Low_1 | 20,781 | 162376 | 41309 | 258008960 | 1539 |
| Low_2 | 42,392 | 421460 | 154957 | 706025723 | 1611 |
| Low_3 | 34,224 | 324587 | 154352 | 535200951 | 1584 |
| Low_4 | 47,768 | 503745 | 65271 | 845208160 | 1616 |
| Low_5 | 29,187 | 267927 | 191955 | 444031405 | 1599 |
| High_1 | 34,884 | 330551 | 60403 | 546207753 | 1593 |
| High_2 | 33,222 | 298031 | 62547 | 485434416 | 1570 |
| High_3 | 18,572 | 150913 | 59961 | 235849817 | 1512 |
| High_4 | 27,568 | 250103 | 164733 | 413241518 | 1594 |
| High_5 | 32,127 | 296928 | 59839 | 486702256 | 1579 |
| Control_1 | 19,545 | 163861 | 79145 | 261507437 | 1550 |
| Control_2 | 23,502 | 209097 | 40255 | 338319404 | 1569 |
| Control_3 | 28,276 | 269744 | 63345 | 445072177 | 1596 |
| Control_4 | 26,291 | 234889 | 61618 | 384602861 | 1589 |
| Control_5 | 17,695 | 143294 | 40606 | 224537265 | 1518 |

**Table S2**. Relative abundance bacterial phyla (>0.5%) in the three experimental groups

| **Taxonomy** | Experimental group | | |
| --- | --- | --- | --- |
|  | **Low** | **High** | **Control** |
| Actinomycetota | 64,83 | 64,23 | 65,97 |
| Pseudomonadota | 31,04 | 31,88 | 29,68 |
| Bacillota | 2,57 | 2,34 | 2,68 |
| Planctomycetota | 0,46 | 0,45 | 0,51 |
| Myxococcota | 0,47 | 0,44 | 0,53 |

**Table S3**. Relative abundance bacterial classes (>0.5%) in the three experimental groups

| **Taxonomy** | Experimental group | | |
| --- | --- | --- | --- |
|  | **Low** | **High** | **Control** |
| Actinomycetes | 59,95 | 59,56 | 60,52 |
| Alphaproteobacteria | 25,13 | 25,92 | 24,38 |
| Betaproteobacteria | 4,24 | 4,33 | 3,70 |
| Thermoleophilia | 2,79 | 2,63 | 3,14 |
| Bacilli | 2,52 | 2,29 | 2,63 |
| Gammaproteobacteria | 1,67 | 1,62 | 1,59 |
| Rubrobacteria | 1,89 | 1,85 | 2,09 |
| Planctomycetia | 0,45 | 0,44 | 0,50 |
| Myxococcia | 0,47 | 0,44 | 0,53 |
| Opitutae | 0,01 | 0,01 | 0,01 |
| Kiritimatiellia | 0,01 | 0,01 | 0,01 |
| Phycisphaerae | 0,01 | 0,01 | 0,01 |

**Table S4**. Relative abundance bacterial genera (>0.5%) in the three experimental groups

| **Taxonomy** | Experimental group | | |
| --- | --- | --- | --- |
|  | **Low** | **High** | **Control** |
| *Streptomyces* | 31,153 | 30,891 | 32,447 |
| *Bradyrhizobium* | 14,189 | 15,062 | 13,458 |
| *Nocardioides* | 8,096 | 7,504 | 8,009 |
| *Starkeya* | 3,766 | 3,672 | 3,801 |
| *Mycobacterium* | 3,861 | 3,855 | 3,715 |
| *Mycolicibacterium* | 3,519 | 3,685 | 3,328 |
| *Methylobacterium* | 1,979 | 1,825 | 1,974 |
| *Streptosporangium* | 1,910 | 1,910 | 1,936 |
| *Baekduia* | 1,872 | 1,834 | 2,077 |
| *Mesorhizobium* | 1,334 | 1,473 | 1,262 |
| *Conexibacter* | 1,204 | 1,128 | 1,317 |
| *Arthrobacter* | 1,045 | 1,061 | 0,914 |
| *Capillimicrobium* | 1,017 | 0,979 | 1,148 |
| *Micromonospora* | 0,831 | 0,903 | 0,829 |
| *Priestia* | 0,781 | 0,747 | 0,791 |
| *Rhodococcus* | 0,850 | 0,859 | 0,784 |
| *Pseudomonas* | 0,353 | 0,261 | 0,244 |
| *Blastococcus* | 0,677 | 0,658 | 0,635 |
| *Sphingomonas* | 0,712 | 0,690 | 0,789 |
| *Actinacidiphila* | 0,583 | 0,778 | 0,564 |
| *Paenibacillus* | 0,567 | 0,422 | 0,598 |
| *Massilia* | 0,589 | 0,605 | 0,562 |
| *Thauera* | 0,636 | 0,721 | 0,523 |
| *Kutzneria* | 0,479 | 0,487 | 0,434 |
| *Peribacillus* | 0,467 | 0,449 | 0,479 |
| *Modestobacter* | 0,495 | 0,497 | 0,480 |
| *Bacillus* | 0,464 | 0,460 | 0,513 |
| *Variovorax* | 0,499 | 0,630 | 0,487 |
| *Salmonella* | 0,266 | 0,344 | 0,307 |
| *Pseudonocardia* | 0,422 | 0,406 | 0,501 |
| *Amycolatopsis* | 0,566 | 0,671 | 0,472 |
| *Kribbella* | 0,441 | 0,464 | 0,406 |
| *Paraconexibacter* | 0,429 | 0,381 | 0,523 |

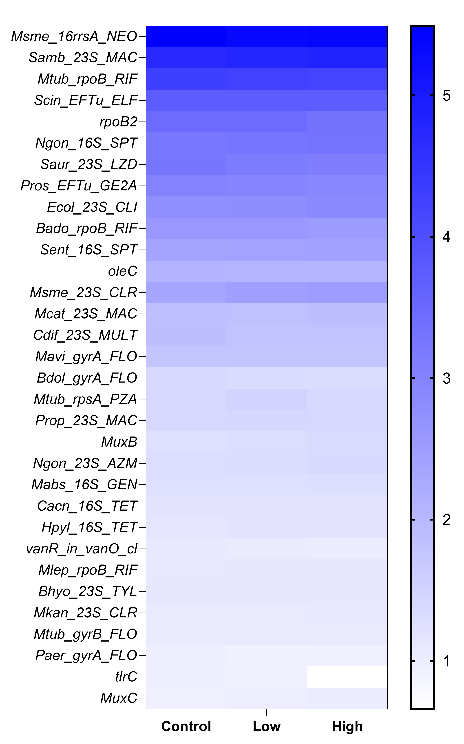

**Figure S1.** ARG profiles and differences between experimental groups. Heatmap showing abundances of 32 top (>1%) ARGs in 3 groups of soil metagenomes.

**Table S5.** The details of potential ARG hosts and their correspondingly harbored ARGs and drug class.

| **Genus** | **Gene** | **Drug class** |
| --- | --- | --- |
| *Actinomarinicola* | Prop_23S_MAC | Macrolide |
| *Afipia* | Nmen_16S_SPT | Aminoglycoside |
|  | Mtub_katG_INH | Isoniazid |
|  | Hpyl_23S_CLR | Macrolide |
|  | Mhom_23S_MAC | Macrolide |
| ***Baekduia*** | novA | Aminocoumarin |
|  | Mtub_rpsL_STR | Aminoglycoside |
|  | Msme_16rrsA_NEO | Aminoglycoside |
|  | Saur_23S_LZD | Aminoglycoside |
|  | Scin_EFTu_ELF | Elfamycin |
|  | Pros_EFTu_GE2A | Elfamycin |
|  | Mtub_gyrA_FLO | Fluoroquinolone |
|  | Paer_gyrA_FLO | Fluoroquinolone |
|  | Bdol_gyrA_FLO | Fluoroquinolone |
|  | Mavi_gyrA_FLO | Fluoroquinolone |
|  | vanR_in_vanO_cl | Glycopeptide |
|  | oleC | Macrolide |
|  | Ctra_23S_MAC | Macrolide |
|  | Cdif_23S_MULT | Multidrug |
|  | Mgal_23S_PLM | Pleuromutilin |
|  | rphA | Rifamycin |
|  | rpoB2 | Rifamycin |
|  | Mtub_rpoB_RIF | Rifamycin |
|  | Bado_rpoB_RIF | Rifamycin |
| ***Bradyrhizobium*** | RbpA | Rifamycin |
| ***Capillimicrobium*** | novA | Aminocoumarin |
|  | Mtub_rpsL_STR | Aminoglycoside |
|  | Msme_16rrsA_NEO | Aminoglycoside |
|  | Saur_23S_LZD | Aminoglycoside |
|  | Scin_EFTu_ELF | Elfamycin |
|  | Pros_EFTu_GE2A | Elfamycin |
|  | Ecol_EFTu_PLV | Elfamycin |
|  | Mtub_gyrA_FLO | Fluoroquinolone |
|  | Paer_gyrA_FLO | Fluoroquinolone |
|  | Bdol_gyrA_FLO | Fluoroquinolone |
|  | vanR_in_vanO_cl | Glycopeptide |
|  | oleC | Macrolide |
|  | Ctra_23S_MAC | Macrolide |
|  | Cdif_23S_MULT | Multidrug |
|  | Mgal_23S_PLM | Pleuromutilin |
|  | rpoB2 | Rifamycin |
|  | Mtub_rpoB_RIF | Rifamycin |
|  | Bado_rpoB_RIF | Rifamycin |
| ***Conexibacter*** | novA | Aminocoumarin |
|  | Mtub_rpsL_STR | Aminoglycoside |
|  | Msme_16rrsA_NEO | Aminoglycoside |
|  | Saur_23S_LZD | Aminoglycoside |
|  | Scin_EFTu_ELF | Elfamycin |
|  | Pros_EFTu_GE2A | Elfamycin |
|  | Mtub_gyrA_FLO | Fluoroquinolone |
|  | Bdol_gyrA_FLO | Fluoroquinolone |
|  | vanR_in_vanO_cl | Glycopeptide |
|  | oleC | Macrolide |
|  | Ctra_23S_MAC | Macrolide |
|  | Cdif_23S_MULT | Multidrug |
|  | Mgal_23S_PLM | Pleuromutilin |
|  | rpoB2 | Rifamycin |
|  | Mtub_rpoB_RIF | Rifamycin |
|  | Bado_rpoB_RIF | Rifamycin |
| *Corynebacterium* | Msme_16rrsA_NEO | Aminoglycoside |
| *Enhydrobacter* | MuxB | Multidrug |
| *Frigoriglobus* | Sent_16S_SPT | Aminoglycoside |
|  | Paer_gyrA_FLO | Fluoroquinolone |
|  | Crei_16rrnS_STR | Aminoglycoside |
|  | Ctra_23S_MAC | Macrolide |
|  | Mhom_23S_MAC | Macrolide |
|  | Cjej_23S_ERY | Macrolide |
|  | Tthe_23S_PLM | Pleuromutilin |
| *Luteitalea* | Pmul_16S_SPT | Aminoglycoside |
|  | Ngon_16S_SPT | Aminoglycoside |
|  | Nmen_16S_SPT | Aminoglycoside |
|  | Sent_16S_SPT | Aminoglycoside |
|  | Crei_16rrnS_STR | Aminoglycoside |
|  | Ecol_23S_CLI | Lincosamide |
|  | Ctra_23S_MAC | Macrolide |
|  | Mhom_23S_MAC | Macrolide |
|  | MexF | Multidrug |
|  | MuxB | Multidrug |
| ***Mesorhizobium*** | RbpA | Rifamycin |
| *Miltoncostaea* | vanR_in_vanO_cl | Glycopeptide |
|  | oleC | Macrolide |
|  | rpoB2 | Rifamycin |
|  | Mlep_rpoB_RIF | Rifamycin |
| ***Nocardioides*** | Mtub_rpsA_PZA | Pyrazine |
| *Paludisphaera* | Ctra_23S_MAC | Macrolide |
| *Pseudolabrys* | Ngon_16S_SPT | Aminoglycoside |
|  | Ecol_23S_CLI | Lincosamide |
|  | Hpyl_23S_CLR | Macrolide |
|  | Mcat_23S_MAC | Macrolide |
|  | ceoB | Multidrug |
|  | MuxB | Multidrug |
|  | rphA | Rifamycin |
|  | Mlep_rpoB_RIF | Rifamycin |
|  | Hpyl_16S_TET | Tetracycline |
|  | Ngon_gyrB_ZOL | Zoliflodacin |
| *Pseudonocardia* | Msme_16rrsA_NEO | Aminoglycoside |
|  | Cdif_23S_MULT | Multidrug |
| *Rhizorhabdus* | novA | Aminocoumarin |
|  | Pros_EFTu_GE2A | Elfamycin |
|  | Mavi_gyrA_FLO | Fluoroquinolone |
|  | vanR_in_vanO_cl | Glycopeptide |
|  | NiCoT NicT Rv2856 | Multidrug |
|  | Mgal_23S_PLM | Pleuromutilin |
|  | rpoB2 | Rifamycin |
|  | Mtub_rpoB_RIF | Rifamycin |
|  | Bado_rpoB_RIF | Rifamycin |
| *Saccharopolyspora* | Cacn_16S_TET | Tetracycline |
| *Singulisphaera* | Sent_16S_SPT | Aminoglycoside |
|  | Scin_EFTu_ELF | Elfamycin |
|  | Pros_EFTu_GE2A | Elfamycin |
|  | Ecol_EFTu_PLV | Elfamycin |
|  | Paer_gyrA_FLO | Fluoroquinolone |
|  | Bdol_gyrA_FLO | Fluoroquinolone |
|  | Mavi_gyrA_FLO | Fluoroquinolone |
|  | vanR_in_vanO_cl | Glycopeptide |
|  | oleC | Macrolide |
|  | Ctra_23S_MAC | Macrolide |
|  | Mhom_23S_MAC | Macrolide |
|  | Cjej_23S_ERY | Macrolide |
|  | Mgal_23S_PLM | Pleuromutilin |
|  | rphA | Rifamycin |
|  | Mtub_rpoB_RIF | Rifamycin |
|  | Bado_rpoB_RIF | Rifamycin |
| *Sphingomonas* | novA | Aminocoumarin |
|  | Scin_EFTu_ELF | Elfamycin |
|  | Pros_EFTu_GE2A | Elfamycin |
|  | Bdol_gyrA_FLO | Fluoroquinolone |
|  | vanR_in_vanO_cl | Glycopeptide |
|  | oleC | Macrolide |
|  | rpoB2 | Rifamycin |
|  | Mtub_rpoB_RIF | Rifamycin |
|  | Bado_rpoB_RIF | Rifamycin |
| *Streptomyces* | Sent_16S_SPT | Aminoglycoside |
|  | Msme_16rrsA_NEO | Aminoglycoside |
|  | Scin_EFTu_ELF | Elfamycin |
|  | Pros_EFTu_GE2A | Elfamycin |
|  | Mtub_gyrA_FLO | Fluoroquinolone |
|  | Paer_gyrA_FLO | Fluoroquinolone |
|  | Bdol_gyrA_FLO | Fluoroquinolone |
|  | Mavi_gyrA_FLO | Fluoroquinolone |
|  | vanR_in_vanO_cl | Glycopeptide |
|  | Ecol_23S_CLI | Lincosamide |
|  | oleC | Macrolide |
|  | Mcat_23S_MAC | Macrolide |
|  | Ctra_23S_MAC | Macrolide |
|  | Mhom_23S_MAC | Macrolide |
|  | Cdif_23S_MULT | Multidrug |
|  | Mgal_23S_PLM | Pleuromutilin |
|  | rphA | Rifamycin |
|  | rpoB2 | Rifamycin |
|  | Mtub_rpoB_RIF | Rifamycin |
|  | Mlep_rpoB_RIF | Rifamycin |
|  | Bado_rpoB_RIF | Rifamycin |
| *Streptosporangium* | Mabs_16S_GEN | Aminoglycoside |
|  | Mtub_rpsL_STR | Aminoglycoside |
|  | Msme_16rrsA_NEO | Aminoglycoside |
|  | Scin_EFTu_ELF | Elfamycin |
|  | Ecol_23S_CLI | Lincosamide |
|  | oleC | Macrolide |
|  | Samb_23S_MAC | Macrolide |
|  | rpoB2 | Rifamycin |
|  | Mtub_rpoB_RIF | Rifamycin |
|  | Bado_rpoB_RIF | Rifamycin |
|  | Cacn_16S_TET | Tetracycline |

Here, only co-occurrence patterns with *ρ* ≥ 0.7 were present.

**Table S6.** Details information of 37 high and medium-quality MAGs detected in this study.

| Experimental group | MAG ID | Classification | Completeness | Contamination | Genome  Size (Mbp) | GC, % | Number of ARGs | Number of BGCs |
| --- | --- | --- | --- | --- | --- | --- | --- | --- |
| Low | MAG_1.5 | c__Gammaproteobacteria | 84.30 | 2.34 | 2.34 | 63.19 | 2 | 7 |
|  | MAG_2.2 | c__Binatia; g__DP-1 | 69.24 | 4.77 | 4.06 | 53.86 | 2 | 5 |
|  | MAG_2.371 | c__Alphaproteobacteria; g__Methylovirgula | 50.00 | 0.00 | 1.6 | 59.50 | 0 | 2 |
|  | MAG_2.76 | c__Alphaproteobacteria; g__Hyphomicrobium_B | 65.00 | 0.86 | 3.22 | 59.44 | 1 | 7 |
|  | MAG_3.17 | c__Alphaproteobacteria; f__Xanthobacteraceae | 50.00 | 6.90 | 3.62 | 63.58 | 1 | 4 |
|  | MAG_3.2 | c__Binatia; g__DP-1 | 62.45 | 3.09 | 3.76 | 54.26 | 1 | 2 |
|  | MAG_3.59 | c__Gemmatimonadetes; g__JACDDX01 | 50.22 | 4.59 | 2.2 | 65.31 | 0 | 1 |
|  | MAG_3.70 | c__Limnocylindria; g__UBA5189 | 51.77 | 7.87 | 2.06 | 70.91 | 1 | 0 |
|  | MAG_3.86 | c__Alphaproteobacteria; g__Methylovirgula | 58.15 | 0.49 | 2.3 | 58.81 | 1 | 0 |
|  | MAG_4.49 | c__Binatia; g__DP-1 | 70.26 | 3.55 | 4.19 | 53.92 | 2 | 2 |
|  | MAG_4.77 | c__Alphaproteobacteria; g__Methylovirgula | 93.78 | 2.25 | 2.84 | 59.73 | 1 | 3 |
|  | MAG_5.32 | c__Alphaproteobacteria; g__Methylovirgula | 85.68 | 0.69 | 2.47 | 59.91 | 1 | 4 |
|  | MAG_5.40 | c__Binatia; g__DP-1 | 70.60 | 3.19 | 3.18 | 54.19 | 0 | 1 |
|  | MAG_5.42 | c__Gammaproteobacteria | 63.40 | 0.34 | 1.87 | 62.62 | 1 | 4 |
| High | MAG_6.12 | c__Alphaproteobacteria; g__Pseudolabrys | 60.16 | 4.31 | 2.73 | 60.65 | 1 | 3 |
|  | MAG_6.14 | c__Binatia; g__DP-1 | 66.71 | 2.28 | 4.21 | 54.20 | 1 | 3 |
|  | MAG_6.58 | c__Gemmatimonadetes; g__JACDDX01 | 64.21 | 1.10 | 2.33 | 65.35 | 0 | 0 |
|  | MAG_7.11 | c__Binatia; g__DP-1 | 74.08 | 1.61 | 3.54 | 54.09 | 1 | 2 |
|  | MAG_7.13 | c__Alphaproteobacteria; g__Pseudolabrys | 54.42 | 3.30 | 2.83 | 60.50 | 1 | 4 |
|  | MAG_9.28 | c__Alphaproteobacteria; g__Pseudolabrys | 56.90 | 1.72 | 3.36 | 60.16 | 0 | 5 |
|  | MAG_9.56 | c__Binatia; g__DP-1 | 64.96 | 2.26 | 3.03 | 54.07 | 1 | 4 |
|  | MAG_9.64 | c__Alphaproteobacteria; g__Methylovirgula | 62.07 | 0.00 | 2.11 | 59.67 | 1 | 3 |
|  | MAG_10.17 | c__Alphaproteobacteria; g__Pseudolabrys | 56.42 | 3.32 | 2.76 | 60.55 | 1 | 4 |
|  | MAG_10.22 | c__Limnocylindria; g__UBA5189 | 54.05 | 7.41 | 1.91 | 71.00 | 3 | 1 |
|  | MAG_10.8 | c__Binatia; g__DP-1 | 70.75 | 3.90 | 3.8 | 54.05 | 0 | 1 |
| Control | MAG_21.26 | c__Thermoleophilia;o__Solirubrobacterales | 62.97 | 5.03 | 1.84 | 68.79 | 1 | 1 |
|  | MAG_21.6 | c__Alphaproteobacteria; g__Pseudolabrys | 52.68 | 3.27 | 2.53 | 60.77 | 1 | 5 |
|  | MAG_22.1 | c__Alphaproteobacteria; g__Methyloceanibacter | 51.03 | 3.45 | 1.78 | 63.92 | 0 | 0 |
|  | MAG_22.19 | c__Binatia; g__DP-1 | 50.33 | 0.65 | 2.39 | 54.43 | 1 | 1 |
|  | MAG_22.35 | c__Alphaproteobacteria; g__Pseudolabrys | 54.47 | 2.59 | 2.87 | 60.58 | 1 | 6 |
|  | MAG_23.14 | c__Alphaproteobacteria; g__Pseudolabrys | 57.68 | 3.90 | 2.94 | 60.45 | 1 | 7 |
|  | MAG_23.7 | c__Binatia; g__DP-1 | 65.66 | 2.17 | 3.31 | 54.14 | 2 | 0 |
|  | MAG_23.8 | c__Limnocylindria; g__UBA5189 | 50.05 | 6.64 | 1.76 | 71.02 | 3 | 2 |
|  | MAG_24.15 | c__Alphaproteobacteria;g__Pseudolabrys | 63.60 | 3.64 | 3.15 | 60.32 | 1 | 5 |
|  | MAG_24.7 | c__Binatia; g__DP-1 | 55.93 | 1.68 | 2.64 | 54.42 | 0 | 0 |
|  | MAG_25.16 | c__Actinomycetes; f__Propionibacteriaceae | 50.36 | 7.25 | 1.93 | 63.39 | 0 | 0 |
|  | MAG_25.2 | c__Thermoleophilia;o__Solirubrobacterales | 55.29 | 8.16 | 1.81 | 69.15 | 0 | 4 |

**Table S7.** Distribution of ARGs across 37 high and medium-quality MAGs from different bacterial classes.

| **ARG** | **Drug class / antibiotic** | Gammaproteobacteria | Binatia | Alphaproteobacteria | Gemmatimonadetes | Limnocylindria | Thermoleophilia | Actinomycetes |
| --- | --- | --- | --- | --- | --- | --- | --- | --- |
| *qacG* | disinfecting agents and antiseptics/ benzalkonium chloride | 1 | 0 | 0 | 0 | 0 | 0 | 0 |
| *rsmA* | Fluoroquinolone-diaminopyrimidine-phenicol /  trimethoprim; chloramphenicol | 2 | 0 | 0 | 0 | 0 | 0 | 0 |
| *adeF* | fluoroquinolone - tetracycline / tetracycline | 0 | 4 | 13 | 0 | 0 | 0 | 0 |
| *vanTG* | glycopeptide / vancomycin | 0 | 7 | 0 | 0 | 0 | 1 | 0 |
| *vanYB* | glycopeptide / vancomycin | 0 | 0 | 0 | 0 | 3 | 0 | 0 |
| *otr(A)S.rim* | tetracycline /oxytetracycline | 0 | 0 | 0 | 0 | 1 | 0 | 0 |
| *vanYG* | glycopeptide / vancomycin | 0 | 0 | 0 | 0 | 1 | 0 | 0 |
| *vanWI* | glycopeptide / vancomycin, teicoplanin | 0 | 0 | 0 | 0 | 1 | 0 | 0 |

**Table S8.** Distribution of BGCs across 31 high and medium quality MAGs.

| Experimental group | MAG ID | Classification | NRPS-like | NRPS-like,RiPP-like | Phosphonate | RRE-containing | RiPP-like | NRPS-like,NRPS | Aryl polyene (APE Vf) | Terpene | NRPS | Terpene (carotenoid) | T3PKS | T1PKS | LAP | Redox-cofactor | Acyl_amino_acids,  hserlactone | RiPP-like | Ranthipeptide | RRE-containing,  lassopeptide | Lanthipeptide class II | NAPAA |
| --- | --- | --- | --- | --- | --- | --- | --- | --- | --- | --- | --- | --- | --- | --- | --- | --- | --- | --- | --- | --- | --- | --- |
| Low | MAG_1.5 | c__Gammaproteobacteria | 1 | 1 | 1 | 1 | 0 | 1 | 1 | 1 | 0 | 0 | 0 | 0 | 0 | 0 | 0 | 0 | 0 | 0 | 0 | 0 |
|  | MAG_2.2 | c__Binatia; g__DP-1 | 0 | 0 | 0 | 1 | 1 | 0 | 0 | 1 | 1 | 1 | 0 | 0 | 0 | 0 | 0 | 0 | 0 | 0 | 0 | 0 |
|  | MAG_2.371 | c__Alphaproteobacteria; g__Methylovirgula | 0 | 0 | 0 | 1 | 0 | 0 | 0 | 1 | 0 | 0 | 0 | 0 | 0 | 0 | 0 | 0 | 0 | 0 | 0 | 0 |
|  | MAG_2.76 | c__Alphaproteobacteria; g__Hyphomicrobium_B | 0 | 0 | 0 | 1 | 0 | 0 | 0 | 1 | 1 | 0 | 1 | 0 | 1 | 1 | 1 | 0 | 0 | 0 | 0 | 0 |
|  | MAG_3.2 | c__Binatia; g__DP-1 | 1 | 0 | 0 | 0 | 0 | 0 | 0 | 1 | 0 | 0 | 0 | 0 | 0 | 0 | 0 | 0 | 0 | 0 | 0 | 0 |
|  | MAG_3.17 | c__Alphaproteobacteria; f__Xanthobacteraceae | 1 | 0 | 0 | 1 | 0 | 0 | 0 | 1 | 0 | 0 | 0 | 0 | 0 | 0 | 0 | 1 | 0 | 0 | 0 | 0 |
|  | MAG_3.59 | c__Gemmatimonadetes; g__JACDDX01 | 0 | 0 | 0 | 0 | 0 | 0 | 0 | 0 | 0 | 0 | 0 | 0 | 0 | 1 | 0 | 0 | 0 | 0 | 0 | 0 |
|  | MAG_4.49 | c__Binatia; g__DP-1 | 0 | 0 | 0 | 1 | 0 | 0 | 0 | 1 | 0 | 0 | 0 | 0 | 0 | 0 | 0 | 0 | 0 | 0 | 0 | 0 |
|  | MAG_4.77 | c__Alphaproteobacteria; g__Methylovirgula | 0 | 0 | 0 | 1 | 0 | 0 | 0 | 1 | 0 | 0 | 0 | 0 | 0 | 1 | 0 | 0 | 0 | 0 | 0 | 0 |
|  | MAG_5.32 | c__Alphaproteobacteria; g__Methylovirgula | 1 | 0 | 0 | 1 | 0 | 0 | 0 | 1 | 0 | 0 | 0 | 0 | 0 | 1 | 0 | 0 | 0 | 0 | 0 | 0 |
|  | MAG_5.40 | c__Binatia; g__DP-1 | 0 | 0 | 0 | 1 | 0 | 0 | 0 | 0 | 0 | 0 | 0 | 0 | 0 | 0 | 0 | 0 | 0 | 0 | 0 | 0 |
|  | MAG_5.42 | c__Gammaproteobacteria | 1 | 0 | 0 | 0 | 0 | 1 | 1 | 0 | 1 | 0 | 0 | 0 | 0 | 0 | 0 | 0 | 0 | 0 | 0 | 0 |
| High | MAG_6.12 | c__Alphaproteobacteria; g__Pseudolabrys | 0 | 0 | 0 | 0 | 0 | 0 | 0 | 1 | 0 | 0 | 0 | 0 | 0 | 1 | 0 | 0 | 1 | 0 | 0 | 0 |
|  | MAG_6.14 | c__Binatia; g__DP-1 | 0 | 0 | 0 | 1 | 0 | 0 | 0 | 1 | 0 | 1 | 0 | 0 | 0 | 0 | 0 | 0 | 0 | 0 | 0 | 0 |
|  | MAG_7.11 | c__Binatia; g__DP-1 | 0 | 0 | 0 | 1 | 0 | 0 | 0 | 1 | 0 | 0 | 0 | 0 | 0 | 0 | 0 | 0 | 0 | 0 | 0 | 0 |
|  | MAG_7.13 | c__Alphaproteobacteria; g__Pseudolabrys | 0 | 0 | 0 | 0 | 0 | 0 | 0 | 1 | 0 | 0 | 1 | 0 | 0 | 1 | 0 | 0 | 1 | 0 | 0 | 0 |
|  | MAG_9.28 | c__Alphaproteobacteria; g__Pseudolabrys | 0 | 0 | 0 | 0 | 0 | 0 | 0 | 1 | 1 | 0 | 1 | 0 | 0 | 0 | 0 | 0 | 1 | 0 | 1 | 0 |
|  | MAG_9.56 | c__Binatia; g__DP-1 | 0 | 0 | 0 | 1 | 1 | 0 | 0 | 1 | 1 | 0 | 0 | 0 | 0 | 0 | 0 | 0 | 0 | 0 | 0 | 0 |
|  | MAG_9.64 | c__Alphaproteobacteria; g__Methylovirgula | 0 | 0 | 0 | 1 | 0 | 0 | 0 | 1 | 0 | 0 | 0 | 0 | 0 | 0 | 0 | 0 | 0 | 0 | 0 | 1 |
|  | MAG_10.17 | c__Alphaproteobacteria; g__Pseudolabrys | 0 | 0 | 0 | 0 | 0 | 0 | 0 | 1 | 0 | 0 | 0 | 1 | 0 | 0 | 1 | 0 | 1 | 0 | 0 | 0 |
|  | MAG_10.22 | c__Limnocylindria; g__UBA5189 | 0 | 0 | 0 | 0 | 0 | 0 | 0 | 1 | 0 | 0 | 0 | 0 | 0 | 0 | 0 | 0 | 0 | 0 | 0 | 0 |
|  | MAG_10.8 | c__Binatia; g__DP-1 | 0 | 0 | 0 | 0 | 0 | 0 | 0 | 1 | 0 | 0 | 0 | 0 | 0 | 0 | 0 | 0 | 0 | 0 | 0 | 0 |
| Control | MAG_21.26 | c__Thermoleophilia;  o__Solirubrobacterales | 0 | 0 | 0 | 0 | 0 | 0 | 0 | 0 | 0 | 0 | 0 | 0 | 0 | 0 | 0 | 1 | 0 | 0 | 0 | 0 |
|  | MAG_21.6 | c__Alphaproteobacteria; g__Pseudolabrys | 0 | 0 | 0 | 0 | 0 | 0 | 0 | 1 | 0 | 0 | 0 | 0 | 1 | 1 | 0 | 0 | 1 | 0 | 1 | 0 |
|  | MAG_22.1 | c__Alphaproteobacteria; g__Methyloceanibacter | 0 | 0 | 0 | 0 | 0 | 0 | 0 | 1 | 0 | 0 | 1 | 0 | 0 | 0 | 0 | 1 | 0 | 0 | 0 | 0 |
|  | MAG_22.19 | c__Binatia; g__DP-1 | 0 | 0 | 0 | 0 | 0 | 0 | 0 | 1 | 0 | 0 | 0 | 0 | 0 | 0 | 0 | 0 | 0 | 0 | 0 | 0 |
|  | MAG_22.35 | c__Alphaproteobacteria; g__Pseudolabrys | 0 | 0 | 0 | 0 | 0 | 0 | 0 | 1 | 0 | 0 | 1 | 1 | 0 | 0 | 0 | 1 | 1 | 0 | 1 | 0 |
|  | MAG_23.14 | c__Alphaproteobacteria; g__Pseudolabrys | 1 | 0 | 0 | 1 | 0 | 0 | 0 | 1 | 0 | 0 | 0 | 0 | 1 | 1 | 0 | 0 | 1 | 1 | 0 | 0 |
|  | MAG_23.8 | c__Limnocylindria; g__UBA5189 | 1 | 0 | 0 | 0 | 0 | 0 | 0 | 1 | 0 | 0 | 0 | 0 | 0 | 0 | 0 | 0 | 0 | 0 | 0 | 0 |
|  | MAG_24.15 | c__Alphaproteobacteria;  g__Pseudolabrys | 1 | 0 | 0 | 0 | 0 | 0 | 0 | 1 | 0 | 0 | 1 | 0 | 0 | 0 | 0 | 0 | 1 | 0 | 1 | 0 |
|  | MAG_25.2 | c__Thermoleophilia;  o__Solirubrobacterales | 0 | 0 | 0 | 1 | 1 | 0 | 0 | 1 | 1 | 0 | 0 | 1 | 0 | 0 | 0 | 1 | 0 | 0 | 0 | 0 |

**Table S9.** Distribution of BGCs across 31 high and medium-quality MAGs from different bacterial classes.

| BGC | Gammaproteobacteria | Binatia | Alphaproteobacteria | Gemmatimonadetes | Limnocylindria | Thermoleophilia |
| --- | --- | --- | --- | --- | --- | --- |
| NRPS-like | 2 | 1 | 4 | 0 | 1 | 0 |
| NRPS-like,RiPP-like | 1 | 0 | 0 | 0 | 0 | 0 |
| Phosphonate | 1 | 0 | 0 | 0 | 0 | 0 |
| RRE-containing | 1 | 6 | 7 | 0 | 0 | 1 |
| RiPP-like | 0 | 2 | 0 | 0 | 0 | 0 |
| NRPS-like,NRPS | 2 | 0 | 0 | 0 | 0 | 0 |
| Aryl polyene (APE Vf) | 2 | 0 | 0 | 0 | 0 | 0 |
| Terpene | 1 | 8 | 15 | 0 | 2 | 1 |
| Non-ribosomal peptide synthetase (NRPS) | 1 | 2 | 2 | 0 | 0 | 1 |
| Terpene (carotenoid) | 0 | 2 | 0 | 0 | 0 | 0 |
| T3PKS | 0 | 0 | 6 | 0 | 0 | 0 |
| T1PKS | 0 | 0 | 2 | 0 | 0 | 0 |
| LAP | 0 | 0 | 3 | 0 | 0 | 0 |
| Redox-cofactor | 0 | 0 | 7 | 1 | 0 | 0 |
| Acyl_amino_acids, hserlactone | 0 | 0 | 2 | 0 | 0 | 0 |
| RiPP-like | 0 | 0 | 3 | 0 | 0 | 2 |
| Ranthipeptide | 0 | 0 | 8 | 0 | 0 | 0 |
| RRE-containing, lassopeptide | 0 | 0 | 1 | 0 | 0 | 0 |
| Lanthipeptide class II | 0 | 0 | 4 | 0 | 0 | 0 |
| Non-alpha poly-amino acids (NAPAA) | 0 | 0 | 1 | 0 | 0 | 0 |
